# TUG protein acts through a disordered region to organize the early secretory pathway

**DOI:** 10.1101/2024.08.09.607373

**Authors:** Anup Parchure, Helen Tejada, Zhiqun Xi, Maohan Su, Omar Julca-Zevallos, Abel R. Alcázar-Román, You Yan, Xinran Liu, Derek Toomre, Jonathan S. Bogan

## Abstract

The endoplasmic reticulum (ER)-Golgi Intermediate Compartment (ERGIC) is a distinct compartment in mammalian cells, which forms in part by homotypic fusion of ER-derived vesicles and gives rise to the cis cisterna of the Golgi ribbon. How the ERGIC is regulated is not well understood. Here we show that the TUG protein is essential to maintain this compartment as a distinct organelle. TUG (UBXN9, Aspscr1) is known to regulate the cell type -specific trafficking of GLUT4 glucose transporters, but its role in more ubiquitous trafficking pathways has not been well characterized. TUG localized to the ERGIC and in fibroblasts its deletion enhanced anterograde flux and increased resorption of ERGIC markers into the cis-Golgi, perturbing membrane homeostasis in the early secretory pathway. TUG knockout cells had a compacted Golgi morphology, and ultrastructural studies revealed dilated cisterna with surrounding small vesicles. A central disordered region in TUG mediated its recruitment to ERGIC membranes, and an amino terminal domain was sufficient to induce oligomerization in cells. In TUG knockout cells, ERGIC-dependent processes such as autophagy are disrupted and model cargoes such as CFTR are missorted. Together, these results reveal a novel, TUG-dependent regulatory mechanism in the early secretory pathway, which modulates ERGIC organization and anterograde trafficking. This function is co-opted by GLUT4 and other proteins that employ an unconventional secretion pathway to the plasma membrane.

## Main

About a quarter of the proteome enters the secretory pathway in mammalian cells. These proteins, including both soluble and membrane-associated secretory proteins, are synthesized at the endoplasmic reticulum (ER). Properly folded proteins are packaged into COPII-coated vesicles at ER exit sites (ERES)^1^. After uncoating, these vesicles are thought to condense with each other, giving rise to the ER-Golgi intermediate compartment (ERGIC). The ERGIC was originally considered as responsible for the long-range traffic of cargoes from peripheral ERES to the centrosome-adjacent Golgi^2, 3^. More recently, using synchronous cargo release, it was demonstrated that ER-to-Golgi trafficking is supported by an intertwined network of tubules, which is dynamic and decorated by both COPII and COPI coats^4^. Thus, an alternate possibility is that the ERGIC may be contiguous with ERES, with segregation of membrane lipids and proteins resulting from effects of membrane curvature, coat proteins, cargo receptors, and other factors. The ERGIC serves as a station for protein sorting, so that both anterograde and retrograde carriers mediate the further trafficking of proteins and membrane lipids from this compartment^5, 6^. In addition, data support the idea that the ERGIC itself undergoes a maturation process to generate the *cis* cisterna of the Golgi complex^7^. How the sorting of various proteins at the ERGIC occurs, and how the ERGIC matures to form the cis Golgi or gives rise to anterograde carriers, are incompletely understood.

In addition to its role in the conventional ER-Golgi secretory pathway, the ERGIC participates in various other cellular functions. The ERGIC may detect improperly folded proteins, and serve as a backup system for ER-associated degradation^5^. The ERGIC also acts in macroautophagy (hereafter termed ‘autophagy’) and serves as a source of membranes for autophagosome biogenesis^8, 9^. Finally, the ERGIC functions in “unconventional” secretion pathways by which membrane proteins are delivered to endosomes or directly to the plasma membrane, bypassing the Golgi complex^10, 11^. Such unconventional secretion pathways are often cell type -specific and mediate the exocytic translocation of various physiologically important membrane proteins. For example, CFTR, a chloride channel that is mutated in cystic fibrosis, is thought to traffic to the plasma membrane at least in part by using such a Golgi-bypass pathway^12, 13^. It remains unknown how particular membrane trafficking pathways at the ERGIC may be adapted in specific cell types to control the exocytic translocation of specialized transmembrane cargoes.

The GLUT4 glucose transporter is also suggested to traffic by an unconventional, Golgi-bypass pathway^14–16^. In fat and muscle cells, insulin stimulates glucose uptake by mobilizing GLUT4 to the cell surface. In cells not stimulated with insulin, GLUT4-containing vesicles are trapped near the ERGIC by the action of TUG (Tether, containing a UBX domain, for GLUT4; also called Aspscr1, UBXN9) proteins^17–21^. Insulin triggers site-specific endoproteolytic cleavage of TUG to liberate these vesicles and to load them onto kinesin motors for translocation to the cell surface^22, 23^. Data imply that the insulin-responsive vesicles form, at least in part, by budding from ERGIC membranes^24^. Because TUG localizes at the ERGIC, it is ideally positioned to capture these vesicles and to sequester them away from the plasma membrane^19^. Upon insulin stimulation, and in adipocytes with shRNA-mediated TUG depletion, the mobilized vesicles fuse directly at the plasma membrane^25^. The precise mechanism by which TUG traps GLUT4-containing vesicles within unstimulated cells remains uncertain. More broadly, GLUT4 expression and TUG cleavage are cell type -specific processes, observed in fat and muscle cells but not in other cell types^20, 22, 26^. TUG itself was discovered >20 years ago and is widely expressed, yet its only well-characterized role has been in the trafficking of GLUT4^17, 27^. To mediate insulin-responsive GLUT4 trafficking, fat and muscle cells may co-opt a more ubiquitous function of TUG. This function is not understood.

Here we identify a more general function for TUG. We show that TUG is critical to maintain the ERGIC as a distinct compartment in the early secretory pathway. Cells lacking TUG exhibit a distorted Golgi morphology, together with accelerated flux of cargoes from the ER to the cis-Golgi. TUG contains two intrinsically disordered regions (IDRs) and can form liquid-like biomolecular condensates *in vitro*. A central IDR is necessary and sufficient for TUG localization to the ERGIC in cells, and an N-terminal domain is sufficient for TUG oligomerization *in trans*. Finally, cellular processes that rely on the ERGIC are disrupted in TUG knockout cells. These data identify TUG as an essential protein to organize the early secretory pathway and imply that this function is co-opted in specialized cell types to mediate unconventional secretion pathways that bypass the Golgi complex.

## Results

### TUG protein localizes to the ERGIC and organizes the early secretory pathway

Previous results show that endogenous TUG protein is present both in punctate structures, colocalized with the ERGIC marker ERGIC53, and diffusely throughout the cytosol^17, 19^. As well, when an extended linker is used, TUG can be tagged at its C-terminus without disrupting its function in 3T3-L1 adipocytes^22^. Therefore, to image TUG protein in cells without fixation or antibody staining, we used a linker to fuse mCherry fluorescent protein at the TUG C-terminus. In TUG knockout (KO) HeLa cells, this TUG-mCherry protein was enriched in punctate structures that colocalized extensively with Emerald-tagged ERGIC53^28^, and in a diffuse pattern throughout the cytosol and nucleus, recapitulating previous results (Figs. 1a, S1a). When Emerald-ERGIC53 was expressed alone, this marker had a more prominent juxtanuclear distribution (Fig. 1b) and it partially overlapped with GM130 and with Sec31A, proteins present at the cis-Golgi and at ERES, respectively (Figs. S1b, S1c). Expression of TUG-mCherry appeared to concentrate ERGIC53 preferentially in peripheral punctate structures. TUG-mCherry was excluded from perinuclear ERGIC53 structures and did not overlap significantly with the Golgi, as shown by co-expression of monomeric Neon green (mNG) tagged GMAP210 (Fig. 1a, Fig. S1d). Syntaxin 12 (Stx12, also called Stx13) is consistently associated with GLUT4 in proteomic studies^29–33^, and is also required for the unconventional secretion of CFTR^13^. We observed partial overlap of TUG-mCherry with GFP-tagged Stx12, (Fig. S1e). These results are expanded upon further below. Together, the data are consistent with previous results and show that TUG localizes at the ERGIC in HeLa cells.

**Figure 1.**
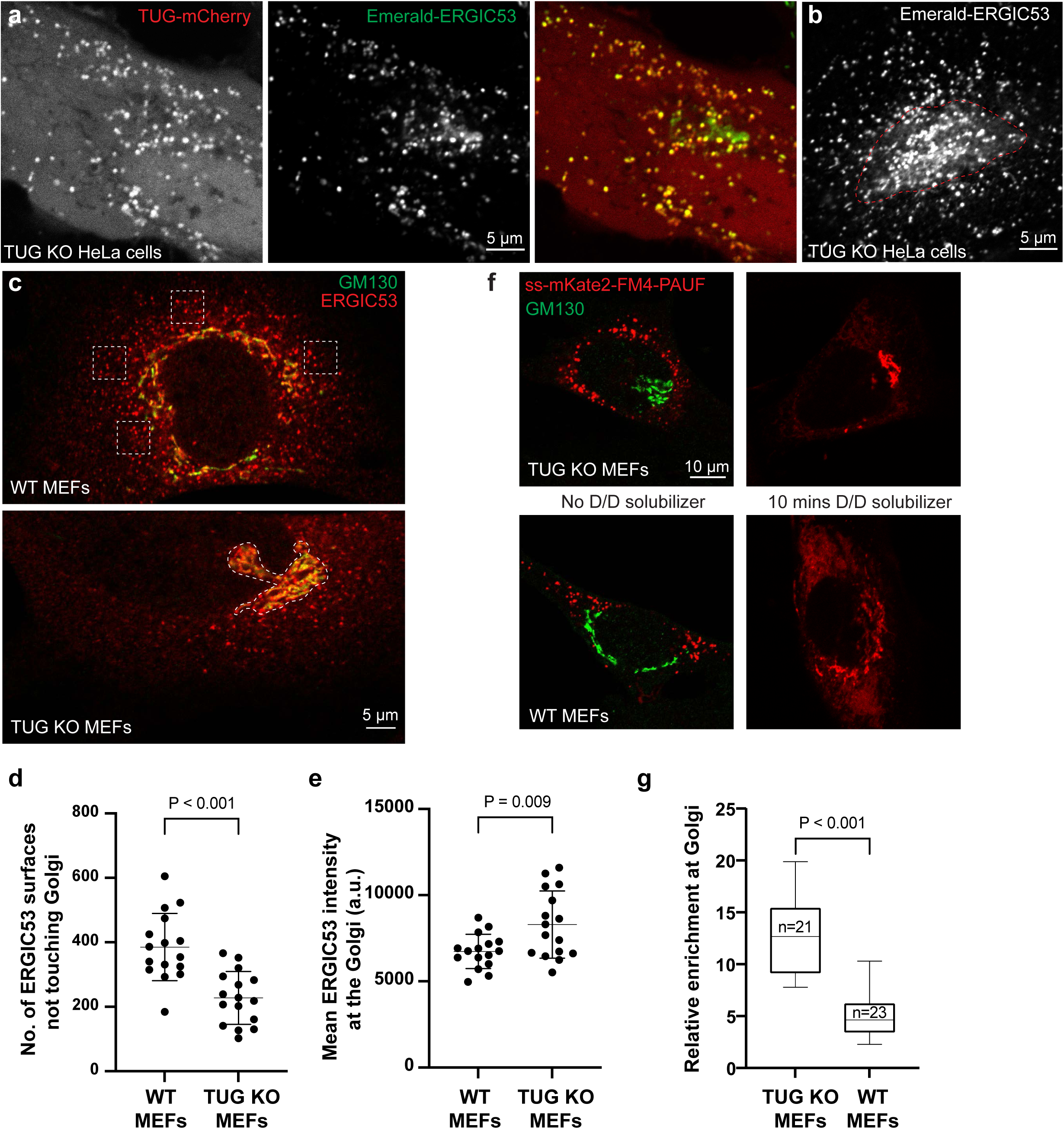
TUG localizes to the ERGIC and regulates anterograde trafficking. a) Images from TUG KO HeLa cells co-transfected with mCherry-tagged TUG (Tug-mCherry; red) and Emerald tagged ERGIC53 (Em-ERGIC53; green). Note the colocalization between TUG and ERGIC53 in peripheral punctate structures. b) Images from TUG KO HeLa cells transfected with Em-ERGIC53 alone shows ERGIC53 distribution in peripheral punctate structures that are in close apposition to the ERES (see Fig. S1c), as well as in a juxtanuclear distribution (red dashed region) that stains positive for the cis-Golgi marker GM130 (see Fig. S1b). c) WT (top) and TUG KO (bottom) MEFs were fixed and stained using antibodies to the ERGIC marker ERGIC53 (green) and the cis-Golgin GM130 (red). Images are a single slice from a confocal stack. The dashed boxes in the top image denote the peripheral punctate staining of ERGIC53, which is devoid of the GM130 signal. The dashed region in the bottom image denotes the Golgi, based on GM130 staining. Note the reduction in peripheral ERGIC53 puncta in TUG KO MEFs, compared to WT control cells, as well as an increased distribution of the ERGIC53 signal at the cis-Golgi. d) Confocal images were segmented in Imaris to generate ERGIC53 and GM130 surfaces. The graph compares independent ERGIC53 surfaces (not touching GM130 surfaces) in WT and TUG KO MEFs, and shows that there is a significant reduction in the numbers of independent ERGIC53 surfaces in TUG KO MEFs. Data from 16 cells are represented as mean +/- standard deviation (s.d.). Statistical analysis was performed using an unpaired two-tailed t test with Welch’s correction (i.e. not assuming equal variances). e) The graph compares average intensities of ERGIC53 staining at the Golgi after collapsing a confocal stack. Data from 16 cells is represented as mean +/- s.d. Statistical analysis was performed using unpaired two-tailed t test. f) Images of TUG KO (top) and WT (bottom) MEFs infected with retroviruses to expresses PAUF tagged with mKate2 and FM4 domains. The presence of FM4 domains results in aggregation of the protein in the ER and is seen distinct from GM130 (green; left) signal in the absence of D/D solubilizer. After addition of D/D-solubilizer, PAUF is released from the ER and traffics in an anterograde manner through the secretory pathway. Cells were fixed 10 minutes after addition of D/D solubilizer and were imaged (right). In the TUG KO MEFs (top), most of the signal is concentrated at the Golgi, but in WT MEFs there is also surrounding ER signal. g) Data from replicates of the experiment shown in (f) were quantified and plotted, and the enrichment of mKate2-tagged PAUF signal at the Golgi is normalized to the signal in the surrounding ER. Data are plotted from at least 20 cells from 2 independent experiments. The box indicates median and quartile ranges, and whiskers indicate the spread of the data from minimum to maximum. Statistical analysis was performed using an unpaired two-tailed t test.

To study effects of TUG knockout, we used murine embryonic fibroblasts (MEFs). We previously generated mice containing a conditional knockout allele of TUG (TUG^fl/fl^)^26^. Immortalized fibroblasts from WT and TUG^fl/fl^ embryos were treated with Cre recombinase to generate control and KO MEFs. Previous data show that TUG depletion in HeLa cells had only a small effect on Golgi morphology, which required brefeldin A washout to elicit a delay in Golgi reformation in siRNA-treated cells^19^. By contrast, the phenotype was much more marked in MEFs, as shown below. We observed that, compared to HeLa cells, MEFs have ∼30-fold greater abundance of TUG protein, which may help the differences in the magnitudes of the phenotypes (Fig. S2a).

In WT MEFs, endogenous ERGIC53 is present in both peripheral punctate and Golgi associated structures (Fig. 1c). In TUG KO MEFs, ERGIC53 was reduced in the periphery, and we observed increased colocalization of ERGIC53 and GM130 staining. To quantify this redistribution, we segmented 3D-confocal stacks to generate ERGIC53 and GM130 surfaces in WT and TUG KO MEFs. We then compared the number of ERGIC53 structures that did not overlap or touch GM130 structures. As shown in Fig. 1d, there were fewer independent ERGIC53 surfaces, not touching GM130, in TUG KO MEFs, compared to WT control cells. Conversely, the intensity of ERGIC53 staining in areas overlapping with GM130 was slightly but significantly increased in TUG KO MEFs, compared with WT MEFs (Fig. 1e). The data support the idea that there is an increased absorption of ERGIC53 into the cis-Golgi in TUG KO MEFs, compared to WT control cells.

The localization of TUG to the ERGIC, together with redistribution of ERGIC53 to the cis-Golgi in TUG KO MEFs, led us to consider whether TUG restrains the anterograde flux of cargo from the ER to the Golgi. That is, it may serve as a brake on the early secretory pathway. To test this idea, we used a model cargo that can be pulse-released from the ER. The soluble cargo protein pancreatic adenocarcinoma upregulated factor (PAUF) has previously been used to study protein secretion^34^. PAUF can be tagged with both a fluorescent protein (mKate2) and FM4 domain repeats and is targeted to the secretory pathway provided that a signal sequence is present at the N-terminus^35, 36^. The FM4 domain repeats then cause PAUF aggregation and retention in the ER and permit the triggered release of PAUF into the secretory pathway upon addition of a solubilizing drug (D/D solubilizer)^37, 38^.

As shown in Fig. 1f, the mKate2-FM4-PAUF reporter aggregated in the ER and did not overlap with GM130 in the absence of D/D solubilizer, both in WT and in TUG KO MEFs. Ten minutes after the addition of solubilizer, most mKate2-FM4-PAUF was concentrated at the cis-Golgi in TUG KO MEFs. At the same time point in WT MEFs, there was some signal at the cis-Golgi, but also much that remained in the ER. By 15 minutes, mKate2-FM4-PAUF was concentrated at the cis-Golgi in WT cells, and the peripheral ER-accumulated protein had dissipated. By 25 minutes after addition of solubilizer, cargo was present in post-Golgi vesicles, both in WT and TUG KO MEFs (Fig. S2c). Quantification of the signal enrichment at the Golgi at 10 minutes after addition of D/D solubilizer confirms a significant increase in TUG KO cells, compared to WT cells (Fig. 1g). The data show that the speed of cargo transport from the ER to the cis-Golgi is greater in TUG KO MEFs, compared to WT control cells. The data support the idea that TUG acts as a brake on flux through the early secretory pathway, which is released in TUG KO cells to accelerate anterograde trafficking.

### TUG deletion alters Golgi morphology

We hypothesized that in cells lacking TUG, altered trafficking at the ERGIC would result in distorted Golgi morphology. As noted above, siRNA-induced TUG depletion in HeLa cells caused only subtle alterations in Golgi morphology, and brefeldin A removal was required to elicit a robust phenotype^19^. Because TUG is present at greater abundance in MEFs, compared to HeLa cells (Fig. S1e), we considered that the phenotype might be more dramatic. Indeed, as assessed by confocal microscopy of two cis-Golgi markers, GM130 and the KDEL receptor (KDELr), the Golgi was much more compacted in TUG KO MEFs, compared to WT control cells (Fig. 2a). This difference was robust, and we quantified it based on GM130 staining in two different ways. First, the Golgi covered much less of the circumference of the nucleus in TUG KO cells, compared to WT MEFs (Fig. 2b). Second, we characterized the outline of the Golgi itself using the circularity function, which relates the perimeter to the area of the Golgi and returns a number closer to 1 when the Golgi is more nearly circular. By this measure as well, TUG KO cells have a more compact Golgi morphology (Fig. 2c). Thus, TUG knockout in MEFs results in a dramatic, compacted Golgi morphology, as assessed by confocal microscopy.

**Figure 2.**
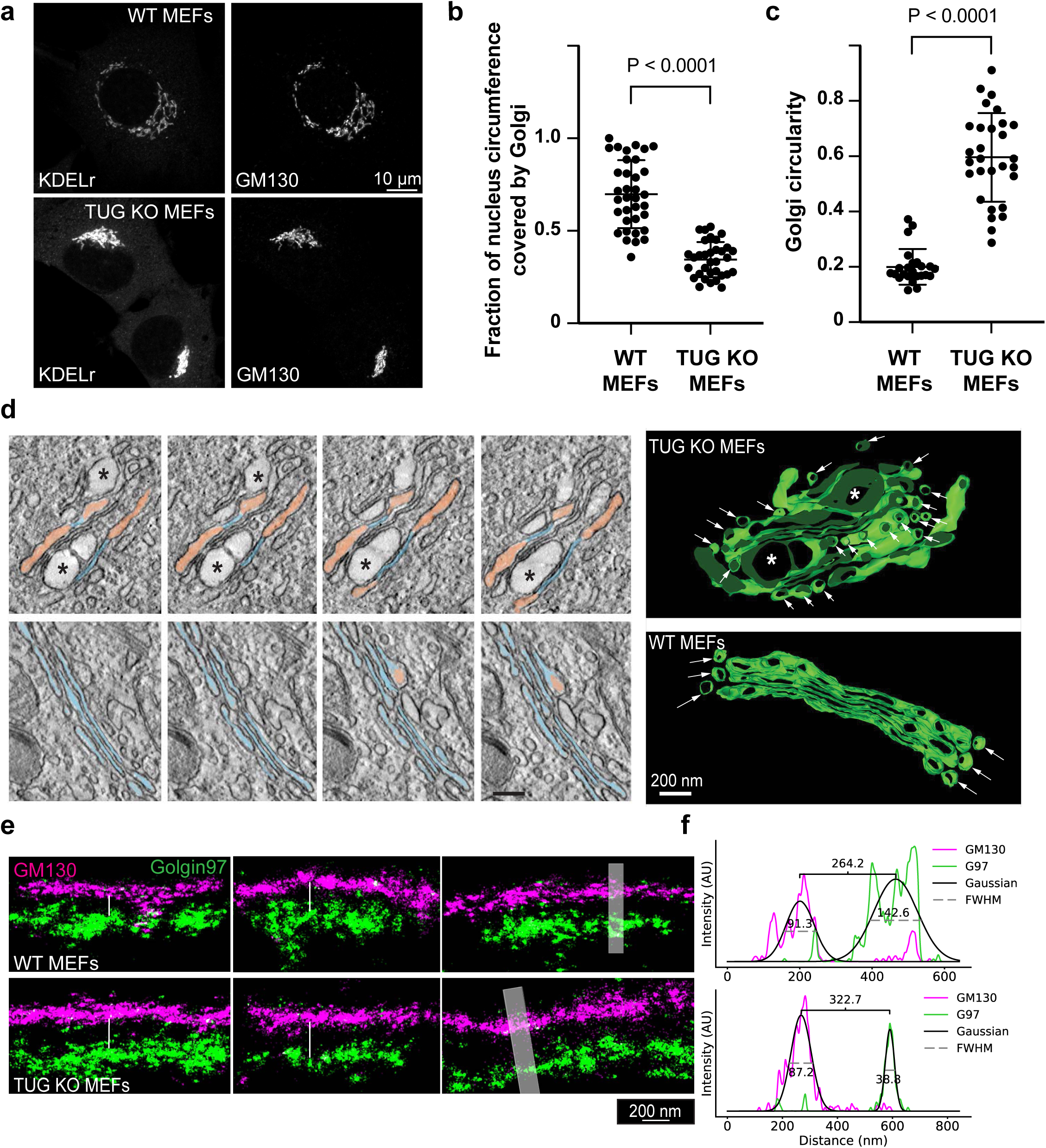
TUG regulates the organization of the Golgi apparatus. a) Images from WT and TUG KO MEFs stained with antibodies to detect KDELr and GM130. Note the compacted Golgi morphology in TUG KO MEFs. b) Quantification of Golgi length along the nuclear circumference. Data are represented as mean +/- s.d. from at least 30 cells. Statistical analysis was performed using a two-tailed t test. c) Quantification of the shape of the Golgi apparatus based on images of GM130 from low magnification images. The compaction of the Golgi was calculated using a circularity function, based on measurements of the perimeter and area of the Golgi complex, as described in the Methods section. Data are represented as mean +/- s.d. from at least 20 cells. Statistical analysis was performed using a two-tailed t test. d) Electron microscopy (EM) was used to image Golgi complexes in WT and TUG KO MEFs. The lumena of narrow, well-stacked cisternae are pseudo-colored in blue, dilated cisternae are in salmon, and ballooned areas are marked by asterisks. At right, EM tomography was used to reconstruct a stack of cross-sectional images through the Golgi complexes of TUG KO MEFs and WT control cells. In addition to the dilated cisterna, an accumulation of small vesicles surrounding the Golgi was observed in TUG KO cells (arrows). e) Representative 4-Pi SMS side view zoomed-in images of Golgi stacks are shown, labeled using antibodies to GM130 (cis-Golgi; magenta) and Golgin97 (trans-Golgi; green). Images from 3 WT MEFs and 3 TUG KO MEFs are shown. The image on the extreme right was used to quantify the separation between the cis and the trans markers. f) Line scan profiles were generated from processed images to obtain the spread between the staining of GM130 and Golgin97. The intensity profile in each channel was fitted to a gaussian and the peak-to-peak distance was used to measure the separation between the cis and the trans markers. In the example shown, this distance was 264 nm in WT MEFs and 323 nm in TUG KO MEFs, so that the increase in the cis-trans distance was ∼60 nm. FWHM represents Full width at half-maximum.

To characterize the ultrastructural alterations in Golgi morphology in TUG KO MEFs, we used electron microscopy (EM) tomography. This revealed that TUG KO MEFs have both dilated Golgi cisterna and abundant small unfused vesicles surrounding the Golgi stacks (Fig. 2d). The dilated cisterna suggested that there might be an increased distance between cis- and trans-Golgi markers in TUG KO MEFs. Therefore, we also used 4pi Single Molecule Switching (SMS) microscopy to image GM130 (cis-Golgi) and Golgin97 (trans-Golgi) in fixed cells (Fig. 2e). As predicted, the distance between these markers was greater in TUG KO MEFs than in WT MEFs, consistent with the morphological changes observed by EM tomography.

### TUG protein forms condensates in solution

Analysis of TUG protein *in silico* using Alphafold^39^ and PONDR^40^ reveals that along with structured domains (UBL1, UBL2 and UBX), TUG protein contains two predicted intrinsically disordered regions (IDRs) (Fig. S3a, b). The larger of these is a central region, IDR1, encompassing residues 183-321 of murine TUG (Fig S3a and Fig. 3a). The second region, IDR2, is at the C-terminus of the protein and includes residues 462-550. IDRs are associated with proteins that form biomolecular condensates and, indeed, TUG is predicted to undergo liquid-like phase separation according to the MolPhase algorithm^41^. To test whether TUG can form condensates *in vitro*, we expressed an mCherry- and His-tagged TUG protein in BL21 *E. coli*, then purified this protein using nickel-NTA resin followed by gel exclusion chromatography. Upon buffer exchange to 125 mM NaCl, representing physiological salt concentration, we did not observe any condensate formation by confocal microscopy. However, condensate formation could be induced by addition of Ficoll 400, a commonly used crowding agent (Fig. S2c). We further confirmed that these TUG condensates behaved like liquids. By time lapse imaging, we observed fusion of two or more droplets in proximity, followed by rapid relaxation of the condensate to a spherical shape (Fig. S2d). Together, these results suggest that upon molecular crowding, purified TUG protein has the ability to form liquid-like biomolecular condensates.

**Figure 3.**
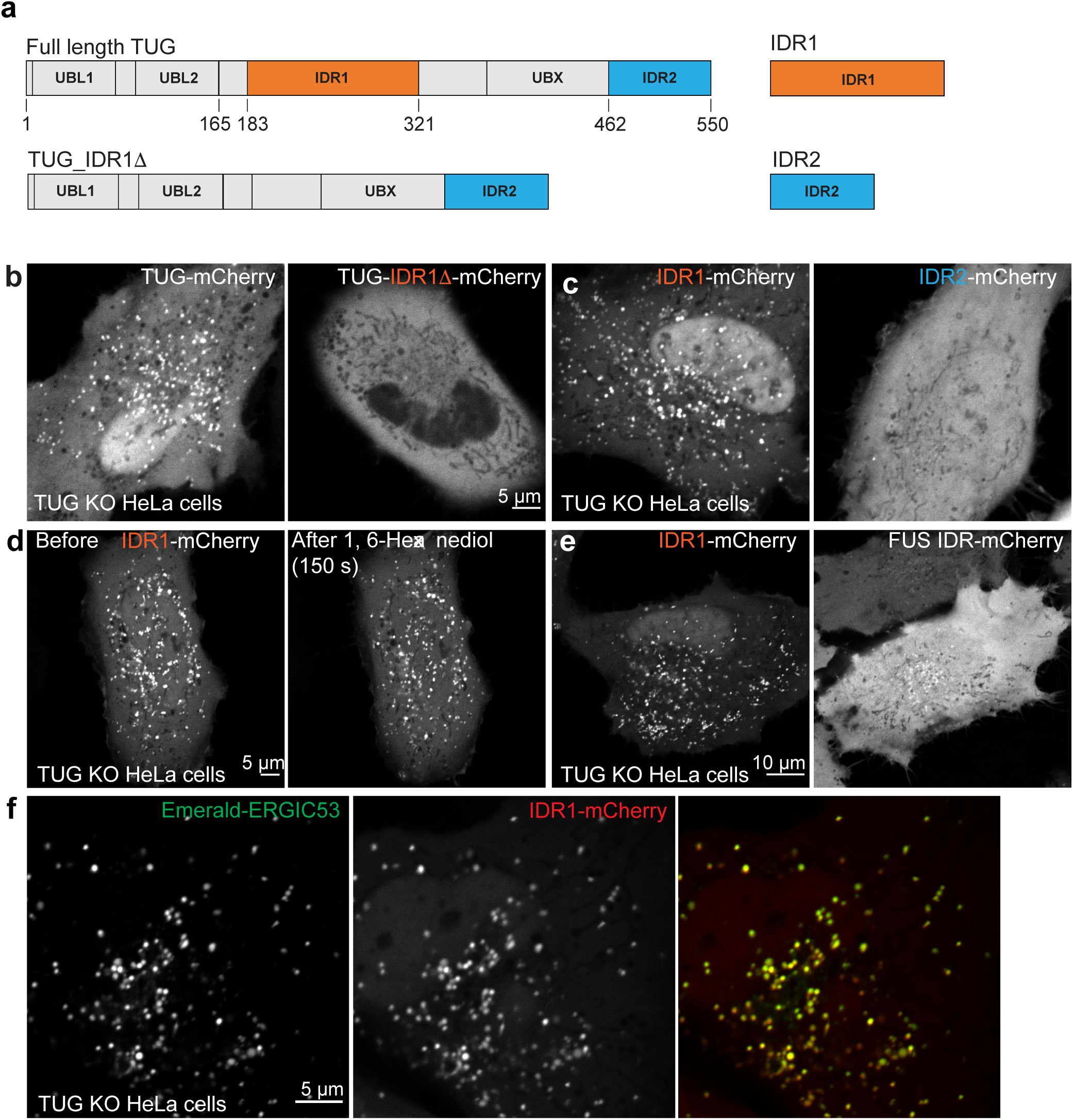
The central disordered region (IDR1) in TUG is necessary and sufficient to mediate TUG localization to the ERGIC. a) Schematic of TUG variants used for expression in TUG KO HeLa cells. All the proteins also contained a C-terminal mCherry tag to aid the visualization of the proteins in living cells. b) Images of TUG KO HeLa cells transfected with full-length TUG (Tug-mCherry; left) or with a variant where the central IDR, IDR1 was deleted (Tug-IDR1Δ-mCherry; right). Although full length TUG forms punctate structures, the mutant is present in a diffuse distribution. Also note that the mutant is excluded from the nucleus. c) Images of TUG KO HeLa cells transfected with IDR1 alone, tagged with mCherry (IDR1-mCherry; left), or with IDR2 alone, also tagged with mCherry (IDR2-mCherry; right). Although IDR1 forms punctate structures in cells, IDR2 is present in a diffuse distribution. d) Images of TUG KO HeLa cells transfected with IDR1-mCherry and treated with 1,6-hexanediol, and imaged before (left) or 150 sec. after 1,6-hexanediol treatment (right). Note that the punctate structures containing IDR1-mCherry continue to be present after 1,6-hexanediol treatment. The structures before and after are not necessarily at the same position, due to shift and refocusing during imaging after drug treatment. e) Images of TUG KO HeLa cells transfected with IDR1-mCherry (left) and mCherry-tagged IDR from FUS protein (right). Note the differences in the distribution of the two IDRs in cells. f) Images from TUG KO HeLa cells co-transfected with IDR1-mCherry (red) and Em-ERGIC53 (green). Note the colocalization of the two proteins in peripheral punctate structures.

### The central IDR mediates TUG localization to the ERGIC

To test whether TUG forms condensates in cells to act in the early secretory pathway, we used truncated proteins and isolated IDRs fused to mCherry. We expressed these proteins in TUG KO HeLa cells and monitored their distribution by confocal microscopy. We reasoned that in cells lacking endogenous TUG, the distribution of ectopically expressed TUG variants would not be influenced by potential effects of oligomerization with endogenous, intact TUG protein. We observed that although full-length TUG had a punctate distribution, proteins lacking the central IDR (TUG-IDR11′-mCherry) were present in a diffuse pattern throughout the cytosol (Fig. 3b). Of note, this protein was also completely excluded from the nucleus. We also performed the converse experiment. We fused IDR1 and IDR2 independently with mCherry and expressed these proteins in TUG KO HeLa cells. Although IDR2 was distributed diffusely, IDR1 formed punctate structures in the cytoplasm and was strongly enriched in the nucleus (Fig. 3c). To confirm that the punctate structures of IDR1 were not due to the mCherry tag or the linker sequence in the fusion protein, we generated another fusion construct by appending a monomeric version of superfolder GFP (sfGFP) and a different linker sequence at the C-terminus of IDR1. Expression of sfGFP-tagged IDR1 in TUG KO HeLa cells resulted in similar punctate distribution (Fig. S4a). We also confirmed that in TUG KO MEFs, the distribution patterns of both IDR1-mCherry and TUG-IDR11′-mCherry were similar to those we observed in HeLa cells (Fig. S4b). These results suggested that the central IDR, IDR1 has the potential to form condensates in cells.

To test whether the punctate structures we observed upon expression of IDR1 are condensates, we treated cells with 1,6-hexanediol, an aliphatic alcohol that has been used to acutely disrupt condensates in cells^42–44^. We did not observe dissolution of IDR1 puncta after hexanediol treatment (Fig. 3d). Due to submicron size of the puncta, it was not possible to photobleach a small region within the condensate and to monitor fluorescence recovery. The spatial distribution of N-terminal IDR from FUS protein, which may form biomolecular condensates^45^, was distinct from the TUG-IDR1 (Fig. 3e). Together with data showing that TUG IDR2 is present in a diffuse pattern (Fig. 3c, above), we conclude that the IDR1 structures are specific and not due to a general feature of expressing intrinsically disordered protein domains in cells.

We wondered whether the IDR1 puncta we observed were isolated structures or associated with particular organelles. We co-expressed mCherry-tagged IDR1 together with Emerald-tagged EGRIC53, using TUG KO HeLa cells as above, and we observed extensive colocalization of the two proteins (Fig. 3d). This result is similar to that using full-length TUG- mCherry (Fig. 1a, above). In addition, mCherry-tagged IDR1 colocalized with a subset of Stx12 structures in TUG KO HeLa cells (Fig. S4c). This result is also similar to data using full-length TUG (Fig. S1e, above). Together, the data indicate that TUG IDR1 is necessary and sufficient to localize TUG to the ERGIC. To test whether TUG and ERGIC53 interact directly, we expressed Emerald-ERGIC53 in HEK293FT cells and immunoprecipitated ERGIC53 using GFP-trap beads. We did not observe copurification of endogenous TUG protein in the immunoprecipitates, suggesting that these proteins do not interact with high affinity (Fig. S4d).

We conclude that the central IDR in TUG is unique and contains localization signals to target TUG to the nucleus and to the ERGIC. Based on current and previous data, both of these targeting signals have important roles in mediating TUG function in cells.

### TUG protein can oligomerize *in trans* via N-terminal ubiquitin-like domains

We next sought to test whether TUG is able to drive the recruitment of ERGIC membranes in cells. Previous studies showed that ectopic localization of golgin proteins, conferred by a mitochondrial targeting signal, is sufficient to capture specific, Golgi-bound vesicles on mitochondria^46^. We wondered if targeting TUG to mitochondria might similarly capture ERGIC membranes at this ectopic location. We used the mitochondrial outer membrane targeting sequence from monoamine oxidase A (MOA) to target TUG-mCherry to mitochondria, and we expressed this TUG-mCherry-mito protein in TUG KO MEFs. Although we did not observe marked recruitment of ERGIC53 to mitochondria, we instead observed dramatic clumping of the mitochondria themselves (Fig. 4a). Control experiments in which only mCherry was targeted to mitochondria showed that the mitochondria had a filamentous structure and were dispersed throughout the cells (Fig. 4b). Thus, the mitochondrial clumping was due to TUG itself, not the mCherry tag or targeting signal. When TUG KO MEFs expressing mitochondrially-targeted TUG were examined by electron microscopy, the mitochondria were stacked against each other, distinct from the dispersed mitochondria observed in control cells (Fig. 4c, d). The data indicate that TUG protein can oligomerize *in trans* when targeted to membrane-bound organelles, and that the affinity of these oligomeric complexes is sufficient to drive mitochondrial clustering.

**Figure 4.**
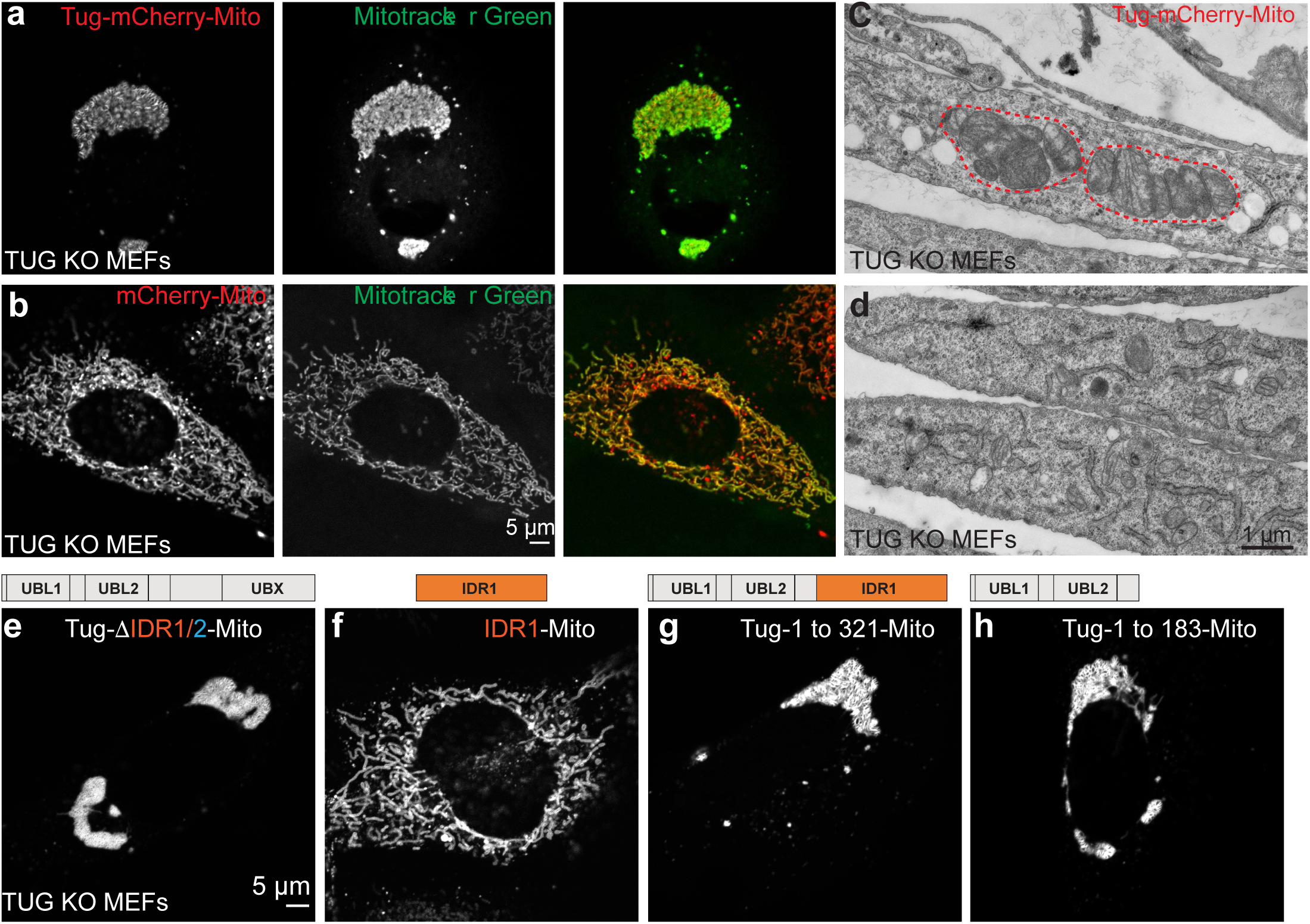
TUG oligomerizes *in trans* via its N-terminal ubiquitin-like domains. a) Images of TUG KO MEFs infected with retroviruses to express mCherry-tagged TUG (red) which was tethered to the mitochondria by the fusion of a transmembrane domain from the protein mitochondrial monoamine oxidase A (Tug-mCherry-Mito). Living cells were also incubated with Mitotracker green (green), which was washed off prior to imaging. Tethering of TUG to the mitochondria results in mitochondrial clumping. b) Images of TUG KO MEFs expressing mCherry tethered to mitochondria (red), together with Mitotracker green (green). Mitochondria are present in a filamentous organization under this condition. c, d) Electron micrographs (negative staining) from TUG KO MEFs expressing Tug-mCherry-Mito (c) or without any exogenous protein expression (control; d). Mitochondria appear stacked when TUG is tethered to the mitochondrial outer membrane by monoamine oxidase A, but are dispersed in control cells. e-h) Images of TUG KO MEFs expressing different truncations of TUG tagged with mCherry and tethered to the mitochondria. Deletion of IDR1 shows that it is not required for clumping of mitochondria (e), and IDR1 alone is not sufficient to clump mitochondria (f). The N-terminal structured UBL1 and UBL2 regions in TUG protein are sufficient to mediate protein-protein interaction *in trans* and thus to clump mitochondria when fused to the transmembrane domain of mitochondrial monoamine oxidase A (g, h).

We reasoned that we could use mitochondrial clustering as an assay to identify regions of TUG that are responsible for its oligomerization. Because we observed TUG condensates *in vitro*, and because condensate formation is typically mediated by interactions among disordered regions, we wondered whether one or both IDRs in TUG would be responsible for the mitochondrial clumping we observed. Accordingly, we expressed a mitochondrially-targeted version of TUG in which both IDRs, IDR1 and IDR2, were deleted. This protein continued to cause clumping of mitochondria, similar to intact TUG (Fig. 4e). Conversely, targeting of IDR1 alone to the mitochondria failed to cause clumping (Fig. 4f). When we expressed a mitochondrially-targeted form of a larger fragment of TUG, containing the UBL1, UBL2, and IDR1 domains (residues 1-321), we again observed clumping of mitochondria (Fig. 4g). Finally, as IDR1 did not appear to mediate this effect, we expressed a mitochondrially-targeted fragment containing residues 1-183. This fragment was sufficient to mediate mitochondrial clumping (Fig. 4h). Of note, this fragment corresponds to the tandem ubiquitin-like (UBL1, UBL2) domains, which are predicted by AlphaFold to extend through residue 173^23^. We conclude that the region of TUG that mediates its oligomerization is the structured N-terminus of the protein.

### TUG regulates autophagy and CFTR trafficking

We hypothesized that cellular functions that rely on the ERGIC might be disrupted in TUG KO MEFs. The ERGIC serves as a source of membranes for autophagosome biogenesis^8, 9^. Thus, we sought to determine if autophagy is impaired in TUG KO cells. We used a Halo-GFP-LC3B reporter protein, which is proteolytically processed during autophagy to generate a protease-resistant Halo ligand-bound fluorescent fragment^47^. The relative abundance of this fragment corresponds to the rate of autophagic flux and can be measured using in-gel fluorescence imaging. We expressed this LC3 reporter using retroviruses in WT and TUG KO MEFs, and isolated cells with similar levels of expression using FACS. We then starved cells to induce autophagy and analyzed processing of the reporter, as diagrammed in Fig. 5a. As shown in Fig. 5b, the abundance of processed LC3 reporter observed by in-gel fluorescence was reduced in TUG KO MEFs, compared to WT control cells. We quantified replicate experiments, which showed that the ratio of processed to unprocessed LC3 reporter was reduced by half in TUG KO cells (Fig. 5c). The ratio of processed to total LC3 reporter, which corresponds to autophagic flux, was also significantly reduced in TUG KO MEFs, compared to WT control MEFs (Fig. 5d). We conclude that TUG deletion causes a reduction in the rate of autophagic flux, consistent with the idea that the ERGIC is regulated by TUG and functions in autophagosome biogenesis.

**Figure 5.**
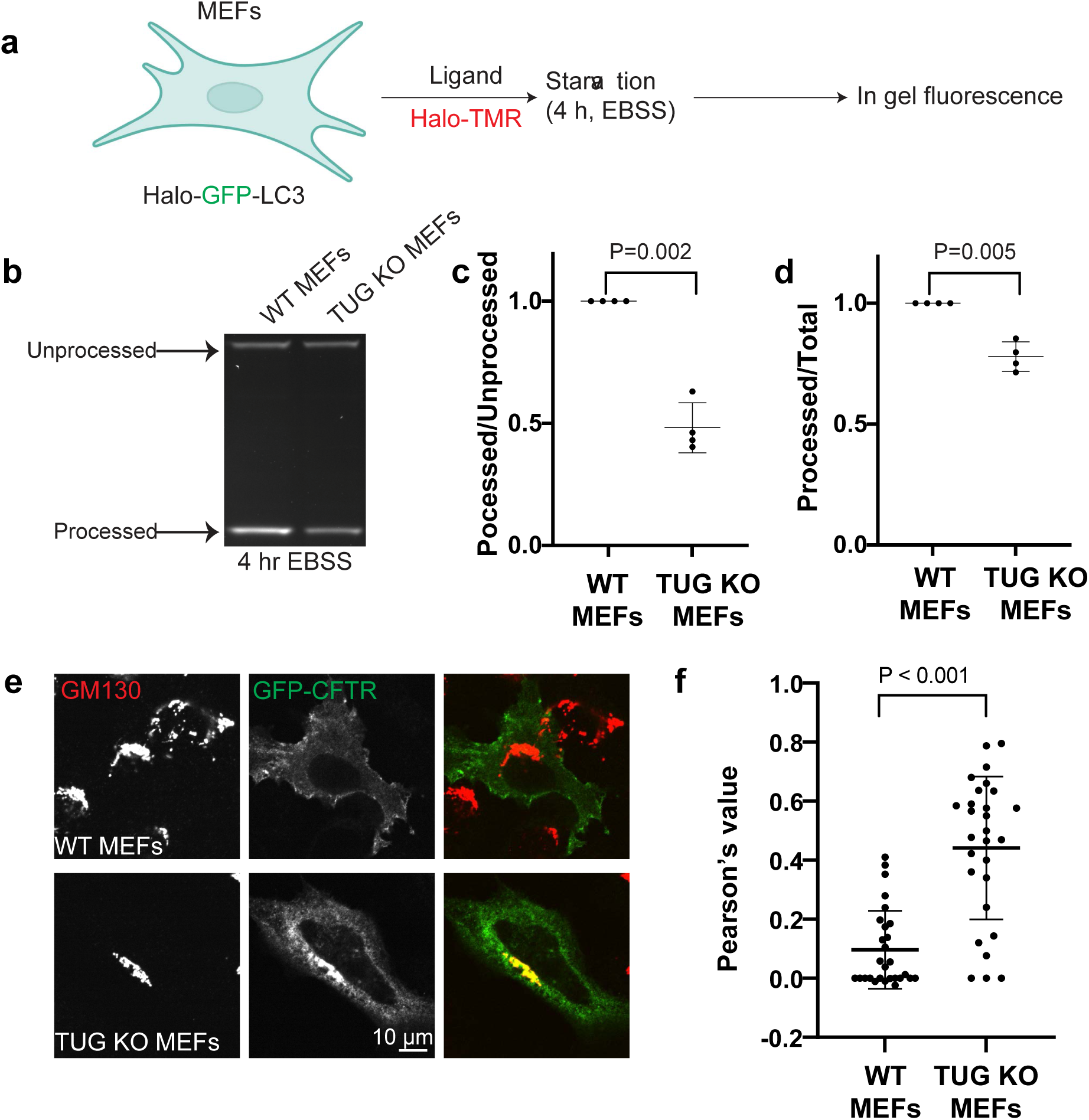
Physiological effects of TUG depletion on autophagy and CFTR trafficking. a) Schematic the autophagy processing assay using Halo-tagged reporters. WT and TUG KO MEFs expressing Halo-and GFP-tagged LC3 were generated using retroviral expression and FACS sorting. Cells stably expressing the reporter construct were incubated with Halo-TMR for 20 minutes. Cells were then incubated in EBSS for 4 hours to induce autophagy, then harvested, lysed and analyzed by SDS-PAGE. Gels were imaged to assess TMR fluorescence. b) A representative in-gel fluorescence image is shown to monitor autophagic flux in WT and TUG KO MEFs. Full length (unprocessed) and Halo-tag (processed) bands are observed in both WT and TUG KO cell lysates. Note the low intensity of the processed band in TUG KO MEFs compared to WT MEFs. c) Quantification of the ratio of band intensities of processed to the unprocessed bands (after background subtraction) from fluorescent images in replicate experiments. The value for WT MEFs is normalized to 1 and the value for TUG KO MEFs is shown in relation to that of WT MEFs from each experiment. Data are represented as mean +/- s.d. from 4 independent experiments. Statistical analysis was performed using a two-tailed t test. d) The quantification of the ratio of band intensities of processed to intact LC3 reporter (processed+unprocessed) in replicate experiments is plotted. Band intensities are calculated after background subtraction from in-gel fluorescent images. The value for WT MEFs is normalized to 1 and the value for TUG KO MEFs is represented in relation to that of WT MEFs from each experiment. Data are represented as mean +/- s.d. from 4 independent experiments. Statistical analysis was performed using a two-tailed t test. e) Images from WT (top) and TUG KO MEFs (bottom) transfected with GFP-tagged CFTR and fixed and stained using GFP booster (green) and an antibody to the cis-Golgin GM130. GFP-CFTR is accumulated at the Golgi in TUG KO MEFs, but not in WT MEFs. f) Pearson’s correlation coefficient was used to measure the overlap of GFP-CFTR and GM130 signal. Data from 28 cells pooled from 3 independent experiments are quantified and presented as mean +/- s.d. Note the increased Pearsons correlation coefficient values for TUG KO MEFs ,which indicates that the CFTR protein accumulates at the cis Golgi upon TUG deletion.

Another function of the ERGIC is to support unconventional secretion pathways, by which particular transmembrane proteins bypass the Golgi and traffic directly to the plasma membrane^11, 48^. We hypothesized that the trafficking of such proteins might be altered in TUG KO cells. To test this idea, we focused on CFTR, which has been shown to traffic, at least in part, by a Golgi-bypass mechanism involving Stx12^12, 13^. We expressed GFP-tagged CFTR^49, 50^ in WT and TUG KO MEFs and monitored the localization of this protein using confocal microscopy. As shown in Fig. 5e, GFP-CFTR protein reached the cell surface in WT cells and did not accumulate in Golgi membranes marked by GM130. In contrast, in TUG KO MEFs, there was marked accumulation of GFP-CFTR at the Golgi, with much less of this protein reaching the plasma membrane. The data support the view that TUG regulates membrane trafficking at the ERGIC, and that the regulated pathways are ones that may be used by proteins that participate in a Golgi-bypass mechanism.

## Discussion

Here we demonstrate that the TUG protein is critical for membrane homeostasis in the early secretory pathway. Our data support the concept that the endoplasmic reticulum (ER) -Golgi intermediate compartment (ERGIC) is a distinct organelle, organized in part through the action of TUG. In cells lacking TUG, the ERGIC is absorbed into the Golgi and the flux of anterograde cargo to the Golgi is accelerated. Increased delivery of cell membranes to the Golgi can account for the distortion of Golgi architecture we observe. As well, cellular functions that depend on the ERGIC, including autophagy and unconventional secretion pathways, are disrupted. Thus, TUG is an essential protein to organize the early secretory pathway, which enables it to serve as a hub for sorting of soluble and transmembrane protein cargoes and for trafficking of membranes.

We propose that TUG acts as a brake in the early secretory pathway, possibly by regulating the biogenesis of anterograde carriers or vesicle fusion at the ERGIC. The increased anterograde cargo flux observed upon TUG deletion may result from relieving this brake. In TUG KO cells, the ERGIC is absorbed into the cis-Golgi, leading to perturbed membrane homeostasis. Thus, a possibility is that TUG KO cells have a compensatory increase in the biogenesis of COPII carriers at the ER. To our knowledge, TUG is first protein described to act as a negative regulator in the early secretory pathway. How cells control the presence of membrane bound organelles in the secretory pathway, and how signaling mechanisms may alter secretory flux to control these organelles, will be interesting avenues for future research.

A central disordered region in TUG is both necessary and sufficient for its recruitment to ERGIC membranes. It remains uncertain precisely how this recruitment occurs. Our data suggest that TUG does not bind with high affinity to ERGIC53, yet it is possible that it interacts with other proteins or membranes present at the ERGIC. As well, our data show that TUG can form biomolecular condensates *in vitro*, but we do not know whether it participates in similar condensates in cells, or whether TUG condensation has a functional role in the organization of the ERGIC. Data suggest other Golgi proteins may form biomolecular condensates, and recent work finds that GM130 co-condenses with RNA to form a phase separated structure linking the Golgi ribbon^51–53^. Other proteins at the early secretory pathway, including TFG and Sec16A, are thought to form biomolecular condensates, which may organize membranes and control cargo flux^54–57^. TUG shares some similarity with TFG, since the disordered C-terminus of TFG mediates its recruitment to the early secretory pathway^58^. Regardless of whether TUG forms condensates *in vivo*, our data show that TUG has the ability to oligomerize *in trans* through its N-terminal ubiquitin-like domains. Possibly, this oligomerization may seed biomolecular condensate formation. Further studies will be required to dissect the function of TUG oligomerization, as the N-terminal UBL domains may also regulate other cellular functions.

Our studies on TUG localization also connect with its role in GLUT4 trafficking. Previous results show that in fat and muscle cells, GLUT4 binds directly to TUG and is retained intracellularly by the action of TUG proteins^18, 26^. Insulin stimulates the endoproteolytic cleavage of TUG to release this trapped GLUT4 and to mobilize it to the plasma membrane^20, 22, 59^. Data support the idea that the N-terminal TUG cleavage product, containing the UBL1 and UBL2 domains, is a ubiquitin-like protein modifier, called TUGUL, which is covalently attached to KIF5B motor proteins. Because this N-terminal product binds directly (and noncovalently) to GLUT4 and IRAP (another transmembrane cargo in the GLUT4 vesicles), its attachment to KIF5B can load these vesicles onto kinesin motors for long-range movement to the cell surface^60, 61^. Data show that these vesicles fuse directly at the plasma membrane^20, 25^. As well, the insulin-responsive vesicles containing GLUT4 are formed, at least in part, by budding at the ERGIC^24^. Thus, our present data add further support to the idea that GLUT4 is translocated to the cell surface by an unconventional, Golgi-bypass pathway^14–16, 19^. It is not clear whether binding of GLUT4 and IRAP to the TUG N-terminal ubiquitin-like domains affects the ability of these domains to oligomerize. Of note, TUG cleavage and insulin-responsive GLUT4 translocation are cell type -specific.

Understanding how TUG regulates traffic at the early secretory pathway will be essential to understanding how this mechanism is adapted to mediate insulin action in fat and muscle cells. Our data raise the possibility that the formation of biomolecular condensates, containing TUG and possibly other components, may be important to trap GLUT4-containing vesicles within unstimulated fat and muscle cells. In nerve terminals, condensates containing the synapsin protein are thought to cluster synaptic vesicles^42, 62, 63^. Synapsin acts with transmembrane proteins present in the vesicles to control vesicle size^64^. Possibly, TUG might act similarly to oligomerize or to form a condensate, and thus to cluster small, GLUT4-containing vesicles. The interaction of GLUT4 and IRAP with such a condensate may enable it to act as a sponge, which could hold the vesicles in an insulin-responsive configuration in unstimulated cells. The formation of biomolecular condensates can be promoted by poly(ADP-ribose)^65, 66^. The main cytosolic poly(ADP-ribose) polymerases, PARP5a and PARP5b, bind to the GLUT4 vesicle cargo protein, IRAP, and may poly(ADP-ribosyl)ate TUG^23, 67^. Possibly, poly(ADP-ribosyl)ation could then promote TUG oligomerization or condensate formation, and thus control the number of insulin-responsive vesicles that are trapped within unstimulated fat and muscle cells.

Previous studies show that TUG binds p97/VCP ATPases, similar to other UBX domain - containing proteins^19, 68–74^. Unlike most other UBX domain-containing proteins, TUG can disassemble the ATPase hexamer *in vitro*, but it is not known if it inhibits p97 in cells. It may serve as an adaptor that recruits ATPase activity to particular targets, similar to other UBX domain-containing proteins. In fat and muscle cells, after insulin-stimulated TUG cleavage, p97 activity is thought to be important to extract the TUG C-terminal cleavage product, which then enters the nucleus and acts with transcriptional regulators to promote oxidative metabolism.

Other effects of p97 action through TUG have not been described. Of note, p97 has been shown to act with other UBX adaptors and with syntaxin 5 to regulate Golgi structure^75–78^. Possibly, TUG may bring p97 activity to ERGIC membranes to control a budding or fusion process. Our data show that a subset of Stx12-positive structures overlaps with TUG puncta in cells. Stx12 has been described to act at endosomes and other sites^79–83^. Our data suggest a role for Stx12 at the ERGIC, but precisely what trafficking pathway it mediates is not known. As noted above, Stx12 was previously described to function in the unconventional secretion of CFTR^13^. Our data show that TUG also regulates CFTR targeting, suggesting that TUG and Stx12 may act together in this pathway. Whether Stx12 functions in GLUT4 trafficking is not known. Further studies will be needed to understand the role of Stx12 at the ERGIC and how it may participate in Golgi-bypass trafficking.

In conclusion, data here show that TUG resides at the ERGIC and controls anterograde flux to the Golgi, and that this function is critical to organize the early secretory pathway. Its recruitment to the ERGIC is mediated by an intrinsically disordered region. At the ERGIC, it can oligomerize and may form biomolecular condensates. These biochemical functions enable to act as a brake on secretory flux from the ER to the Golgi. In addition, the function of TUG to organize ERGIC membranes supports cellular functions such as autophagy and unconventional secretion pathways. Understanding how TUG acts with other machinery to control membrane dynamics will thus have broad implications for understanding cell physiology and a range of human disease.

### Materials and methods Cloning and Constructs

For cloning, all PCR amplifications were done using Phusion High-Fidelity DNA polymerase, which was obtained from Thermo Fischer Scientific. Final vectors were generated using Gibson assembly (New England Biolabs; NEB). Restriction digestions were carried out using enzymes from NEB. All reactions were carried out using manufacturer’s protocols. Plasmids were sequenced using the Yale Keck DNA Sequencing Core facility.

To generate the full-length TUG protein tagged with mCherry at the C-terminus, we used TUG with a linker sequence containing a TEV cleavage site followed by an AviTag (BirA biotinylation site), described previously^22^. PCR amplification of the TUG sequence and linker was carried out using a priming site in the pBICD2 vector^14, 17, 84^. mCherry was amplified using the RINS1 construct (gift from Dr. Dmytro Yushchenko (Addgene plasmid # 107290; http://n2t.net/addgene:107290; RRID: Addgene_107290). The two fragments were then fused using Gibson assembly, and cloned into the double digested (EcoR1, Not1) pB retroviral expression vector^17^. All other truncations of TUG tagged to mCherry were generated using full length protein as a template. To generate TUG IDR1 tagged to sfGFP, sfGFP fragment was amplified using a Chromogranin B tagged to sfGFP construct^85^, which was a gift from Dr. Julia von Blume. To generate mCherry-tagged TUG variants that were tethered to the mitochondrial outer membrane, a gene block (Integrated DNA Technologies; IDT) was synthesized to encode the transmembrane domain from the mitochondrial monoamine oxidase A, which was then used as a fragment in Gibson assembly.

To generate the GMAP210 tagged to mNeon green (mNG), plasmids were obtained from Dr. James Rothman were used as templates to amplify the coding regions for GMAP210 and mNG, which were then assembled into the pB vector using Gibson assembly. Cloning of the FUS IDR tagged to mCherry was achieved by PCR amplifying a fragment containing the 214 residues corresponding to the FUS IDR from the pcDNA 3.2-FUS-1-526aa-V5, a gift from Dr. Aaron Gitler (Addgene plasmid # 29609; http://n2t.net/addgene:29609; RRID: Addgene_29609) and then inserted into pB vector along with a C-terminal mCherry tag and a linker which was the same used for cloning of IDR1 from TUG.

The GFP-CFTR plasmid was previously described^49, 50^ and was a gift from Dr. William Guggino.

For *in vitro* experiments to express mCherry and 6X His tagged TUG protein, 6X His and mCherry were appended at the N-terminus of murine TUG protein separated by a TEV protease cleavage site which also serves as a linker sequence. Fragments were PCR amplified and fused with the double digested pET-15B vector (BamH1 and Nco1) using Gibson assembly.

### Cell culture

HEK293FT (Thermo Fisher Scientific), HeLa cells (Catalog number CCL-2; ATCC) and MEFs were cultured in high glucose DMEM Glutmax (Gibco; 10569044) supplemented with 10% EquaFETAL bioequivalent serum (Atlas Biologicals; EF-0500-A), 100 U/ml penicillin, 100 µg/ml streptomycin and 0.25 μg/mL of Gibco Amphotericin and 2.5 μg/mL plasmocin (Invivogen) (complete medium). Cells were maintained at 37°C in presence of 5% carbon dioxide.

To make TUG knockout HeLa cells, CRISPR-Cas9 genome editing was performed essentially as described^86^. Annealed guide RNA oligonucleotides were designed with the help of the CRISPR design tool (http://crispr.mit.edu), cloned into the Bbs1-digested PX459 V2.0 vector, and transformed into Stbl3-competent *E. coli*. After sequencing to confirm successful cloning, 1.0 µg PX459 V2.0 plasmid containing guide RNA was transfected into 100,000 HeLa cells per well in a six-well plate. Cells were selected with 2 μg/ml puromycin for 2 days to kill untransfected cells. After subsequent replating at single cell density, KO clonal cell lines were identified by Western blotting and confirmed by sequencing of PCR-amplified genomic DNA. The oligos to make the gRNAs to knockout TUG were 5’-caccgCGTGTACACGCAGACTGGGG-3’ and 5’-aaacCCCCAGTCTGCGTGTACACGc-3’. The primers used for PCR to verify knockout were 5’- TGATGGTTTCTTTCCTCTCCTC-3’ and 5’- GGACAGCAGATTTTCCAGTTG-3’.

To generate TUG knockout MEFs, primary cultures of murine embryonic fibroblasts from control mice or mice homozygous for a floxed TUG allele, TUG^fl/fl^, were isolated at embryonic day 13.5, using methods described previously^87^. The TUG^fl/fl^ mice were described previously^26^. Mice were backcrossed to C57BL/6J for several generations prior to isolation of MEFs. Control and floxed MEFs were treated with Ad5CMVCre (Ad-Cre), an adenovirus containing Cre recombinase, which was purchased from the Gene Transfer Vector Core at the University of Iowa. Controls included TUG^fl/fl^ cells not treated with Ad-Cre and WT cells exposed to Ad-Cre. Immortalized cells were immortalized by multiple passaging.

### Transfection

Transfection of HeLa cells was done using FuGENE HD transfection reagent (Promega Corp.) as per the manufacturer’s protocol. Briefly, cells were plated in glass bottom imaging dishes from Cellvis (D35-14-1.5-N) and transfection was done at approximately 50% confluency. For co-transfection of pmEM-ERGIC53 (gift from Dr. Ke Xu (Addgene plasmid # 170717; http://n2t.net/addgene:170717; RRID: Addgene_170717)) or GFP-Stx12 (gift from Dr. Derek Toomre) together with Tug-mCherry or IDR1-mCherry, the ratio of the two DNAs was 1:5 (with the mCherry-tagged constructs, which were in the pB retrovirus vector, being used in higher amounts). In all transfections, 2.5 µl of transfection reagent was used for 1 µg of DNA, and transfection complexes were formed by incubation in Opti-MEM (Gibco) for 20 minutes, following which these were added to cells in a dropwise fashion. The amounts of transfection reagent used was scaled according to the final amount of DNA. Cells were maintained in high glucose DMEM Glutamax complete medium without antibiotics for 48 hours prior to imaging.

For imaging experiments using transfected MEFs, cells (WT or TUG KO, ∼6500 cells) were seeded in 35 mm glass-bottom dishes (Mattek P35G-1.5-14-C) the day before transfection. MEFs were transfected with 1 μg GFP-CFTR^49^ using 2.5 μl FuGENE HD transfection reagent (Promega Corp.).

### Antibody staining

The same protocol was utilized for staining of MEFs and HeLa cells in all experiments, except for those involving imaging of GFP-CFTR together with GM130. Cells were fixed using 4% paraformaldehyde (PFA; Electron Microscopy Sciences) for 20 minutes. PFA solution was made in 1X PHEM buffer (60mM PIPES, 25mM HEPES, 10mM EGTA, and 4mM MgSO4·7H_2_0). Cells were then permeabilized for 5 minutes in PHEM buffer containing 0.3% NP-40 and 0.05% Triton X-100. After permeabilization, cells were blocked in PHEM buffer containing 0.05% NP-40, 0.05% Triton X-100 and 5% normal goat serum (Jackson Immunoresearch). Primary antibodies (overnight incubation) and secondary antibodies (1 hour incubation) were diluted in blocking buffer. Cells were finally washed in 1X phosphate buffered saline (PBS; Gibco) and stored in PBS at 4°C before imaging.

To image samples using 4pi SMS microscopy WT and TUG KO MEFs were seeded on 30mm diameter No. 1.5H round coverslips (Thorlabs) and grew for 1-2 days before fixation by 4% PFA for 15 mins and permeabilization by 0.3% NP40 + 0.05%TX-100 for 3 mins. Cells were processed as described above, except the secondary antibodies were incubated for 2 hours at room temperature. After antibody incubation, samples were post-fixed in 3% PFA + 0.1% GA for 10 min and stored in PBS at 4 °C.

To monitor the localization of GFP-CFTR in WT and TUG KO MEFs, cells were fixed two days after transfection using 10% neutral buffered formalin solution (Sigma) for 10 minutes. After fixing, cells were washed with PBS and permeabilized with 0.1% Triton X-100 for 10 minutes. Cells were then blocked in 5% bovine serum albumin (BSA) in PBS with 0.1% Tween 20 (PBST) for 30 minutes. Following blocking, cells were incubated with the primary antibody GM130 (BD Biosciences) in blocking buffer (5% BSA/PBST) for 1 hour at room temperature (RT). Samples were then washed three times with PBST and incubated with the secondary antibody, Atto488-labeled GFP-booster (Proteintech; gba488), and Hoechst 33342 in blocking buffer for 30 minutes at RT. After three washes with PBS, samples were stored in PBS at 4°C before imaging.

### Western blotting

For western blotting, cell lysates were prepared in cold 1% NP40 buffer containing cOmplete protease inhibitor cocktail (Roche). Samples were lysed on ice for 30 minutes followed by centrifugation at 4°C at 20000 × g for 10 minutes. Pellets were discarded, and cell lysate supernatants were stored at -20°C till further use. Protein estimation was done using Bradford assay (Biorad). Samples were boiled at 95°C for 5 minutes in NuPAGE LDS sample buffer (Invitrogen) containing 3.75 % beta-mercaptoethanol (BME). Samples were electrophoresed using 4%-12% gradient NuPAGE gels (Invitrogen) and NuPAGE MOPS SDS running buffer (Invitrogen). Proteins were transferred from the gels to a nitrocellulose membrane (Biorad) using wet blot system (Invitrogen) and NuPAGE transfer buffer (Invitrogen) containing 10% methanol at 10 volts for 90 minutes. After transfer to membranes, they were incubated with 5% milk made in PBS containing 0.1% Tween20 (PBST) for at least one hour. Membranes were incubated with specific primary (overnight incubation) and HRP-coupled secondary antibodies (one hour incubation) and proteins were detected using chemiluminescence (Pierce ECL; Thermo Fischer Scientific) and imaged on ImageQuant LAS400 (Amersham).

For re-probing the membranes using a different primary antibody, membranes were stripped using the Restore PLUS western blot stripping buffer (Thermo Fischer Scientific) for 15 minutes at room temperature. The membranes were washed in PBST and blocked and re-probed using a different antibody and processes further as described above.

### Electron microscopy

Cells cultured in 10 cm dishes were fixed in 2.5% glutaraldehyde in 0.1 M sodium cacodylate buffer (pH 7.4) for 1 hour at room temperature. After rinsing with buffer, they were scraped in 1% gelatin and spun down in 2% agar to form pellets. Samples were post-fixed in 1% osmium tetroxide for 1 hour, dehydrated in a series of ethanol up to 100%, then infiltrated and embedded in Embed 812 medium (Electron Microscopy Sciences). The blocks were cured in 60°C oven overnight. Thin sections (60 nm) were cut using a Leica ultramicrotome (UC7) and post-stained with 2% uranyl acetate and lead citrate. Sections were examined with a FEI Tecnai transmission electron microscope at 80 kV accelerating voltage, and digital images were recorded with an Olympus Morada CCD camera and iTEM imaging software.

For electron microscopy tomography imaging, 250 nm thick sections were cut using a Leica ultramicrotome, collected on formvar/carbon-coated copper grids, and stained with 2% aqueous uranyl acetate followed by lead citrate. 10 nm PGA gold particles were placed on both sides of the grids as fiducial markers before imaging. The tilt-series (single-axis) were collected using a FEI Tecnai F20 TEM at the accelerating voltage of 200 kV; the tilting range was from -60° to 60° in 1° increments. A FEI Eagle CCD camera (4k x 4k) and SerialEM software were used to collect datasets. Image alignment and 3D reconstruction were performed using IMOD software^88^ and manual tracing of membrane contours.

### 4Pi-SMS imaging

Two-color 4Pi-SMS imaging was done on a custom-build microscope^89^. Sample mounting, image acquisition, and data processing were described as in a previous publication^90^, with modifications to imaging protocol involving changes in acquisition speed and drift correction.

### Retrovirus production

Retroviruses were generated using HEK293FT cells. HEK293FT cells were plated on poly-Lysine (Millipore Sigma) -coated 10 cm^2^ dishes and grown in high glucose DMEM Glutamax complete medium. When the cells were 80% confluent, they were transfected with a plasmid cocktail containing the retroviral vector (containing the gene of interest) and the packaging plasmid, pCL-Eco^91^, using Lipofectamine 2000 (Invitrogen) according to manufacturer’s protocol. 5 µg of DNA for each of the plasmids and 40 µl of Lipofectamine 2000 was used for transfection. On the next day, the medium was replaced with fresh complete medium. 48 hours after transfection, media containing virus particles was collected, and passed through a 0.45 µm filter and stored at 4°C. HEK293FT cells were again supplemented fresh complete medium for one more cycle of virus collection. 72 hours after transfection, media containing virus particles was again collected, as above. The 48 h and 72 h collections were pooled and either used immediately for infection or aliquoted and stored at -80°C.

### ER to Golgi transport assays using mKate2-tagged PAUF

WT and TUG KO MEFs (10,000 cells) were plated on glass bottom imaging dishes from Cellvis (D35-14-1.5-N). The next day, cells were infected with retroviruses to drive the expression of mKate2- and FM4-tagged PAUF using 8ug/ml polybrene (Millipore Sigma). The retroviral expression plasmid, pCX4-ss-mKate2-FM4-PAUF, was a gift from Dr. Yuichi Wakana. Virus containing medium was removed and replaced with high glucose DMEM Glutamax complete medium after 24 hours of infection. Two days after infection, cells were incubated with 1 µM D/D-solubilizer (Takara; 635054) in high glucose DMEM Glutamax complete medium for different time intervals at 37°C, then fixed at RT with 4% PFA in PHEM buffer for 20 minutes. Fixed cells were stained using GM130 antibody (BD Biosciences; 610822), Atto 594 conjugated RFP booster (Proteintech; rba594) to amplify the mKate2 signal and Alexa 488 labeled goat-anti mouse secondary antibody to detect GM130. To monitor the ER retained pool, cells were fixed and processed without addition of the D/D-solubilizer as described above. Imaging was carried out on Zeiss 880 using 63x/1.4 oil objective at room temperature.

### Autophagy flux assays

WT and TUG KO MEFs were infected with retroviruses to drive the expression of Halo- and GFP-tagged LC3. The retroviral expression plasmid was a gift from Dr. Thomas Melia. Cells stably expressing similar amounts of the proteins were FACS sorted using GFP fluorescence. A day before the assay, 0.5 million cells were plated in a 6 well plate. The next day, cells were incubated with Halo-TMR (100nM; Promega) for 20 minutes. Cells were washed twice in PBS and then incubated in Earle’s Balanced Salt Solution (EBSS; Gibco) for 4 hours at 37°C to induce autophagy. For control samples, after the TMR-Halo incubation, cells were washed twice in PBS and harvested as described below. Cells were scraped in ice cold PBS. Cells were spun down for 2 minutes at 2000 × g at 4°C. The supernatant was discarded, and the cell pellet was lysed in 1% NP40 buffer containing cOmplete protease inhibitor cocktail (Roche). Cells were lysed on ice for 30 minutes, then lysates were centrifuged at 20000 × g for 10 minutes at 4°C. Pellets were discarded and cell lysate supernatants were stored at -20°C till further use. Protein content was determined from cell lysates using a Bradford protein assay. Equal amounts of each cell lysate sample were heated in NuPAGE LDS sample buffer (Invitrogen) containing BME for 5 minutes at 95°C. Samples were electrophoresed on 4%-12% gradient NuPAGE gels (Invitrogen) using NuPAGE MPOS SDS running buffer (Invitrogen). After separation of proteins, the gels were transferred to MilliQ water and imaged on ChemiDoc MP Imaging system (Biorad). Autophagic flux was quantified by measuring the band intensities of the processed and the unprocessed bands and represented as a ratio of processed to total (processed+unprocessed) for each condition

### Immunoprecipitation

One million HEK293FT cells were plated in a 6 well plate which was coated with poly-Lysine. Cells were transfected with 1 µg of pmEM-ERGIC53 and 2.5 µl of Lipofectamine 2000. Following day, cells were immunoprecipitated using GFP trap agarose beads (Proteintech; gta) according to manufacturer’s protocols. Briefly cells were lysed in 0.5% NP40 containing buffer and cells were spun down at > 20000 × g for 10 minutes at 4°C. The cell lysate supernatant fraction was then incubated with GFP trap agarose beads which were washed and equilibrated in the dilution buffer (10mM Tris, 150mM NaCl, 0.5mM EDTA) and then incubated with the diluted cell lysate and rotated end-over-end for 1 hour at 4°C. 50 µl of diluted cell lysate was kept aside as input fraction before incubation with the beads. After 1 hour incubation, beads were washed three times and resuspended in 2X NuPAGE LDS sample buffer with BME and boiled at 95°C for 5 mins. Samples were then spun down at 20000 × g for 10 minutes and the supernatant was analyzed by SDS-PAGE.

### Antisera

Antibodies used for immunofluorescence were as follows: GM130 (BD Biosciences; 610822), Sec31A (BD Biosciences; 612350), ERGIC53 (Sigma; E1031), Golgin 97 (Proteintech; 12640-1-AP), Atto 488 conjugated GFP booster (Proteintech; gba488), Atto 595 conjugated RFP booster (Proteintech; rba594). Antibody against KDELr was a kind gift from Dr. Jim Rothman’s laboratory. Fluorophore conjugated secondary antibodies (Alexa 488 and Alexa 568) were obtained from Invitrogen. For 4Pi SMS imaging VHH anti-mouse CF660C (Biotium) and VHH anti-rabbit AF647 (Jackson Immunoresearch) were used. For western blotting, antibodies directed to GFP (Proteintech; P42212) and GAPDH (Millipore Sigma; MAB374) were commercially obtained. The TUG antibody was described previously^17, 21^ and is directed to the TUG C-terminal peptide, which is identical in mice and humans. HRP-conjugated secondary antibodies were obtained from Millipore Sigma.

### Live cell imaging

Prior to imaging living cells for the purposes of monitoring the distribution of proteins, cells were incubated in phenol red free DMEM (Gibco; 21063029) containing 10% EquaFETAL bioequivalent serum. Cells were imaged rapidly on Zeiss 880 using 63x/1.4 oil objective at room temperature. To image mitochondria, cells were labeled with 200 nM Mitotracker green FM (Cell Signaling Technology) for 10 minutes in phenol red free DMEM containing 10% EquaFETAL bioequivalent serum. Cells were washed twice using this medium before imaging on a Zeiss 880 using 63x/1.4 oil objective at room temperature. To monitor the sensitivity of IDR1 puncta to 1,6-Hexanediol, cells were imaged at 37°C in presence of 5% carbon dioxide on a Zeiss 880 using 63x/1.4 oil objective. Cells were imaged without the addition of 1,6-Hexanediol (Millipore Sigma; 240117), then an equal volume of medium containing 10% 1,6-Hexanediol was added to the dishes so as to attain a final concentration of 5% for 150 seconds. Cells were then reimaged as above.

### GFP-CFTR localization assay

WT and TUG KO MEFs were fixed and stained as described above and imaged using an Andor Dragonfly Spinning Disc Confocal Microscope (Oxford Instruments) equipped with 405-, 488-, and 640-nm laser lines, 60x UPLSAPO 1.4NA silicone oil objective, and Sona sCMOS camera with a 6.5 μm pixel size.

### Image analysis

To quantify Golgi morphology, WT and TUG KO MEFs were labeled using antibodies to GM130 and KDELr. Images were acquired on a Zeiss 880 confocal microscope, and z-stacks were collapsed to draw regions outlining Golgi staining. Images were analyzed using FIJI (ImageJ) software. To measure the fraction of the nucleus circumference that was covered by the Golgi, the length of the nuclear perimeter adjacent to the Golgi was measured, as was the full nucleus circumference. The ratio of the length of the Golgi along the nucleus, divided by the length (circumference) of the nucleus, was quantified and plotted. To measure compaction of the Golgi, the area and perimeter of the Golgi were measured based on GM130 staining, and circularity formula (x = perimeter^2^ / 4 π area) was applied.

To quantify ERGIC53 surfaces, confocal images of MEFs stained using GM130 and ERGIC53 antibodies were segmented in Imaris after local background subtraction. Information was extracted so as to obtain the distance of each ERGIC53 surface from a Golgi surface. The ERGIC53 surfaces that were not touching any of the Golgi surfaces were considered as independent ERGIC53 surfaces. The number of such independent surfaces was measured in each cell. Data collected on a cell-by-cell basis were plotted and analyzed.

To monitor mean ERGIC53 intensity at the cis-Golgi, confocal stacks were collapsed. Regions of interest (ROIs) that represent the Golgi area were generated from GM130 images by using the Analyze Particles function in FIJI (ImageJ). ROIs were then overlaid on the ERGIC53 channel to compute mean ERGIC53 intensity. ROIs that covered largest contiguous Golgi signal were used for analysis.

To quantify relative enrichment of mKate2 tagged PAUF upon pulse release, ROIs were drawn in the region corresponding the Golgi apparatus to obtain mean signal intensity from the Golgi, which was then normalized to the surrounding signal from the ER. Care was taken to exclude the nucleus which does not contain any signal. For each cell signal enrichment at the Golgi was represented as ratio of the signal at the Golgi to that divided by the signal in the surrounding ER region.

All 4Pi-SMS images were rendered using Point Splatting mode (10 nm particle size) with Vutara SRX 7.0.06 software (Bruker, Germany). Line-scan profiles were generated by custom code in Fiji-ImageJ and plotted and curve-fitted by custom code in Python 3.

To quantify the differences in localization of GFP-CFTR in WT and TUG KO MEFs, Pearson’s correlations on images were analyzed and quantified using the ImageJ colocalization module.

To quantify signals from images of western blots and in gel fluorescence, images were subjected to background subtraction. ROIs were drawn around the bands of interest and integrated signal intensity was quantified using ImageJ.

### Protein expression, purification of His-tagged proteins

Plasmid coding 6xHis- and mCherry-tagged TUG were transformed into *E. coli* BL21 cells. Bacteria were grown in LB medium with 100 µg/ml ampicillin first in 20 ml overnight at 37°C and then was used to inoculate 2 L LB medium with 100 µg/ml ampicillin for 4 hours at 37°C. Cells were then shifted to 16°C for 10 minutes and were induced by addition of 0.1 mM IPTG and incubated overnight at 16°C. The culture was harvested and resuspended in protein purification buffer (25 mM Tris–HCl (pH 7.4), 500 mM NaCl, 5% glycerol, and 1 mM DTT) containing cOmplete protease inhibitor cocktail (Roche), then lysed using a high-pressure homogenizer. The crude lysate was centrifuged at 30,000 rpm for 45 min at 4°C in a Ti-70 rotor. Supernatant was applied to a prepacked column with Ni-NTA resin (Qiagen), which was equilibrated in protein purification buffer also containing 5 mM imidazole. Protein was eluted after washing the column with buffer containing up to 20 mM imidazole, then eluted with 250 mM imidazole. Eluted fractions were pooled together and concentrated before further purifying using gel filtration using a Superdex 200 size-exclusion column (GE Healthcare) and equilibrated in the buffer with a composition 25 mM Tris–HCl (pH 7.4), 500 mM NaCl, 5% glycerol, and 1 mM DTT. Protein was stored at -80°C after flash freezing in liquid nitrogen.

### Condensate formation assays

To monitor condensate formation, protein was thawed and buffer exchanged to assay buffer (25 mM Tris, 125mM NaCl and 1mM DTT) using Amicon ultra centrifugal filters with a 30kDa cutoff. Protein was diluted to the final desired concentration in the assay buffer, either in presence or absence of Ficoll 400 (Sigma; F2637), and incubated for 5 minutes in PCR tubes. The solution was then plated on to glass bottom imaging dishes from Cellvis (D35-14-1.5-N) and condensates were imaged on Zeiss 880 using a 63x/1.4 oil objective. To monitor fusion of two or more condensates, a continuous time lapse image series was recorded.

## Supplementary Figure Legends

**Supplementary Figure S1:**
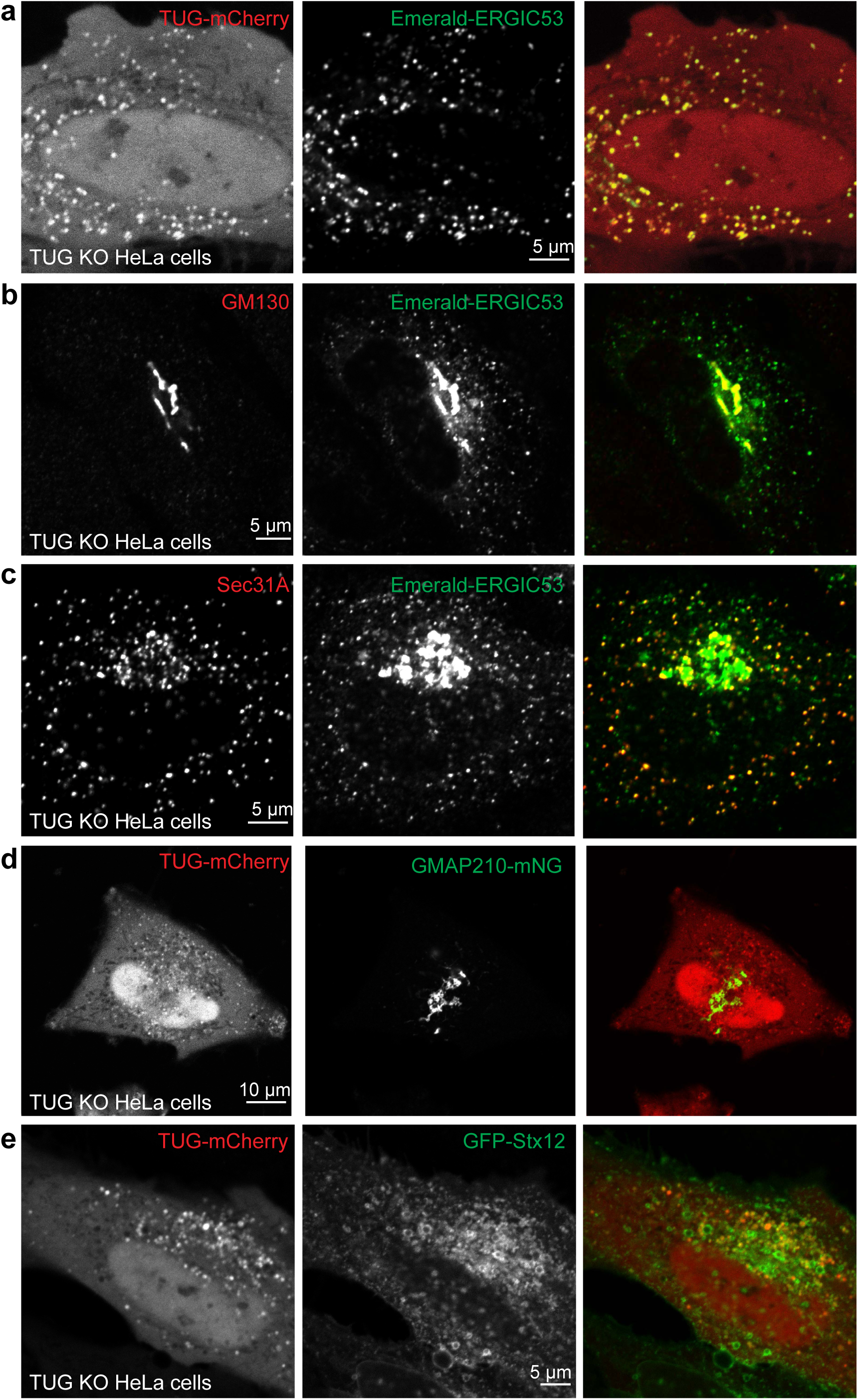
a) An additional set of representative images from TUG KO HeLa cells co-transfected with TUG-mCherry (red) and Em-ERGIC53 (green) is shown to demonstrate the colocalization of these proteins in peripheral punctate structures. b, c) Images of TUG KO HeLa cells transfected with Em-ERGIC53 and fixed and stained with GFP nanobody (green) to amplify the ERGIC53 signal and with antibodies against GM130 (red, b) or Sec31 (red, c) to observe the localization of these proteins at the Golgi and at ERES, respectively. d) Images from TUG KO HeLa cells co-transfected with Tug-mCherry (red) and monomeric Neon Green (mNG) -tagged GMAP210. The images show that the TUG signals are exclusive and do not overlap with the GMAP210 signal. e) Images of TUG KO HeLa cells co-transfected with Tug-mCherry (red) and GFP-tagged Syntaxin 12 (GFP-Stx12, green). Although Stx12 populates structures that are morphologically distinct, a subset of Stx12-positive structures overlaps with the TUG puncta.

**Supplementary Figure S2:**
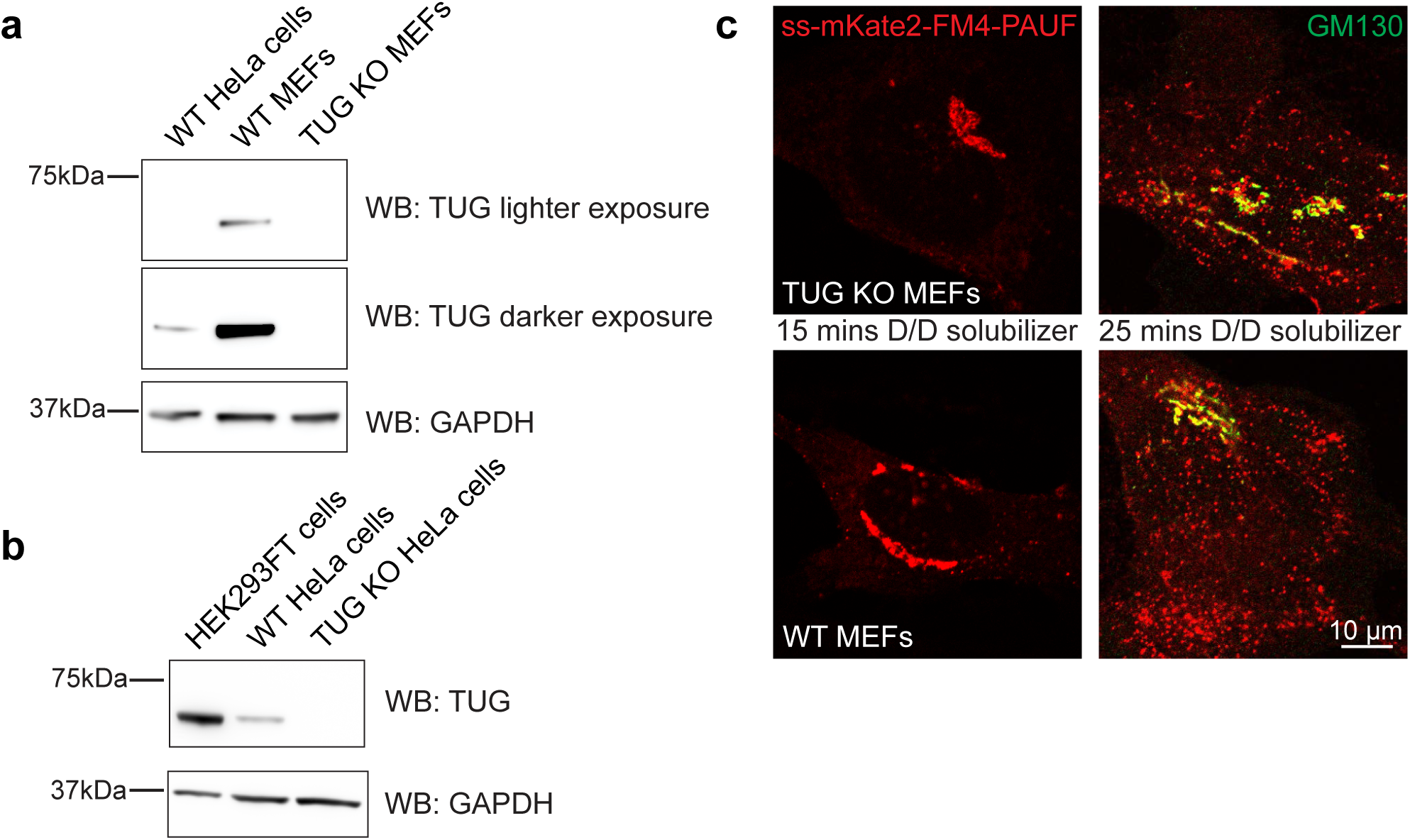
a) Western blot probed using anti-TUG antibody to monitor the relative abundance of TUG protein in WT HeLa cells and WT MEFs, as well as in TUG KO MEFs. In lighter exposure (top), the band is only visible in WT MEFs. However, upon darker exposure (middle), the band is visualized in WT HeLa cells, but no signal is seen from TUG KO MEFs. The bottom panel represents probing the same blot using anti-GAPDH antibody as a loading control. From the western blot, it is evident that HeLa cells express significantly lower levels of TUG abundance, compared to WT MEFs. There is a complete absence of protein expression in TUG KO MEFs, as expected. b) Western blot (top) probed using anti-TUG antibody to monitor the relative abundance of TUG protein in HEK293FT cells, compared to that in WT and TUG KO HeLa cells. The bottom panel represents probing the same blot using anti-GAPDH antibody, as a loading control. Note the absence of protein in TUG KO HeLa cells, but also the lower abundance of the protein in WT HeLa cells as compared to HEK293FT cells. c) Images of ss-mKate2-FM4-PAUF are shown at 15 and at 25 min. after addition of D/D solublizer. In the images taken at 15 min., most PAUF is present at the Golgi, both in TUG KO MEFs and in WT control MEFs. At 25 min., PAUF is present in post-Golgi vesicles in both cell lines.

**Supplementary Figure S3:**
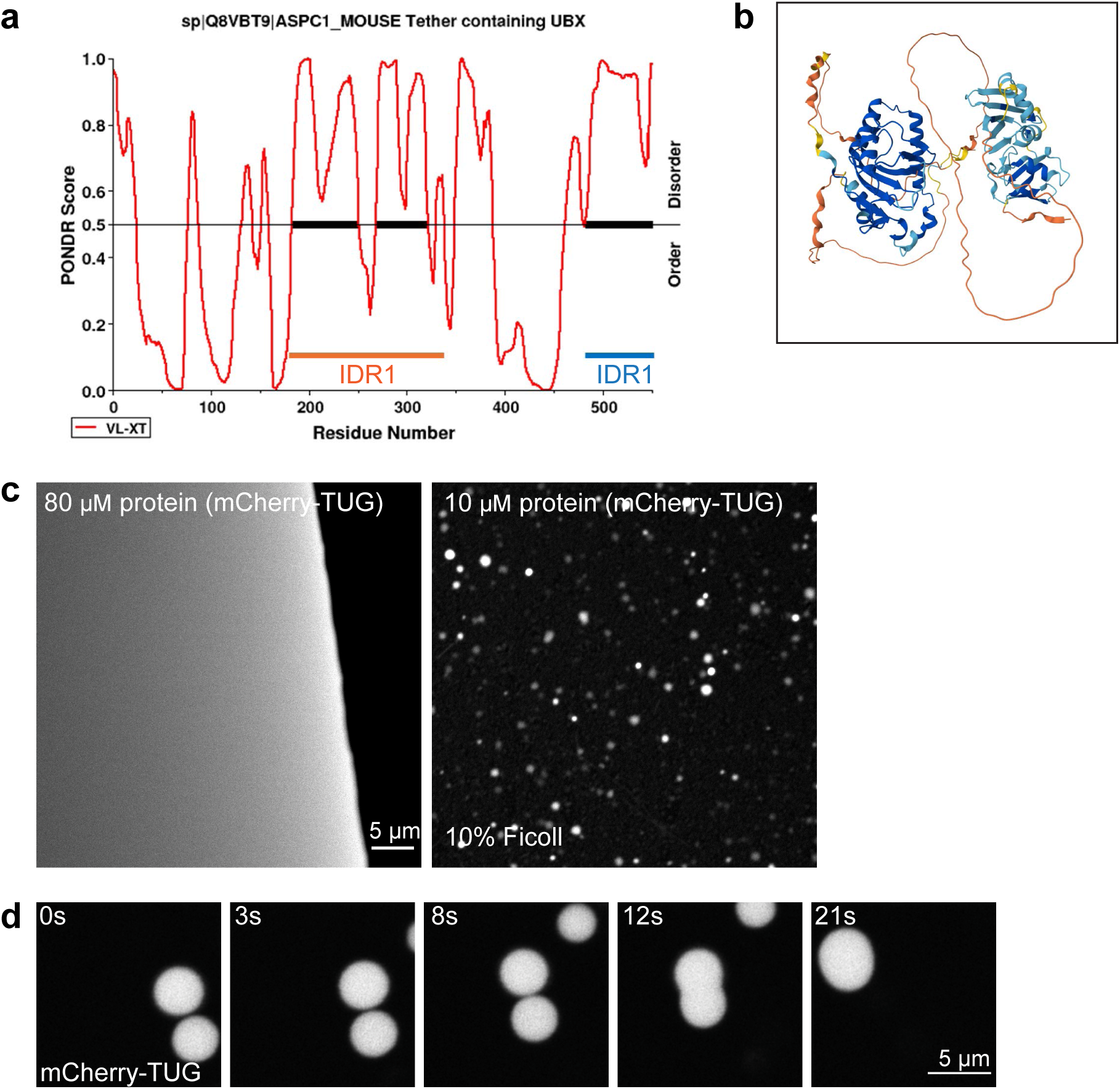
a) Predictions of disordered regions in murine TUG protein were generated using PONDR. TUG protein is characterized by the presence of an internal disordered region (IDR1, residues 183-321, orange) and a smaller disordered region at the C-terminus (IDR2, residues 462-550, blue), when analyzed using the VL-XT algorithm. b) Alpha fold prediction of the murine TUG protein also shows the presence of disordered regions, interspersed between structured regions. c) Purified mCherry tagged TUG protein (80 µM) does not form condensates in phase separation buffer containing 125 mM NaCl (left). Yet, in presence of 10% Ficoll 400, condensates are seen at 10 µM protein concentration. d) A set of images extracted from time lapse imaging demonstrate the coalescence of two condensates in proximity, followed by relaxation to a spherical shape, which is reflective of a liquid-like behavior.

**Supplementary Figure S4:**
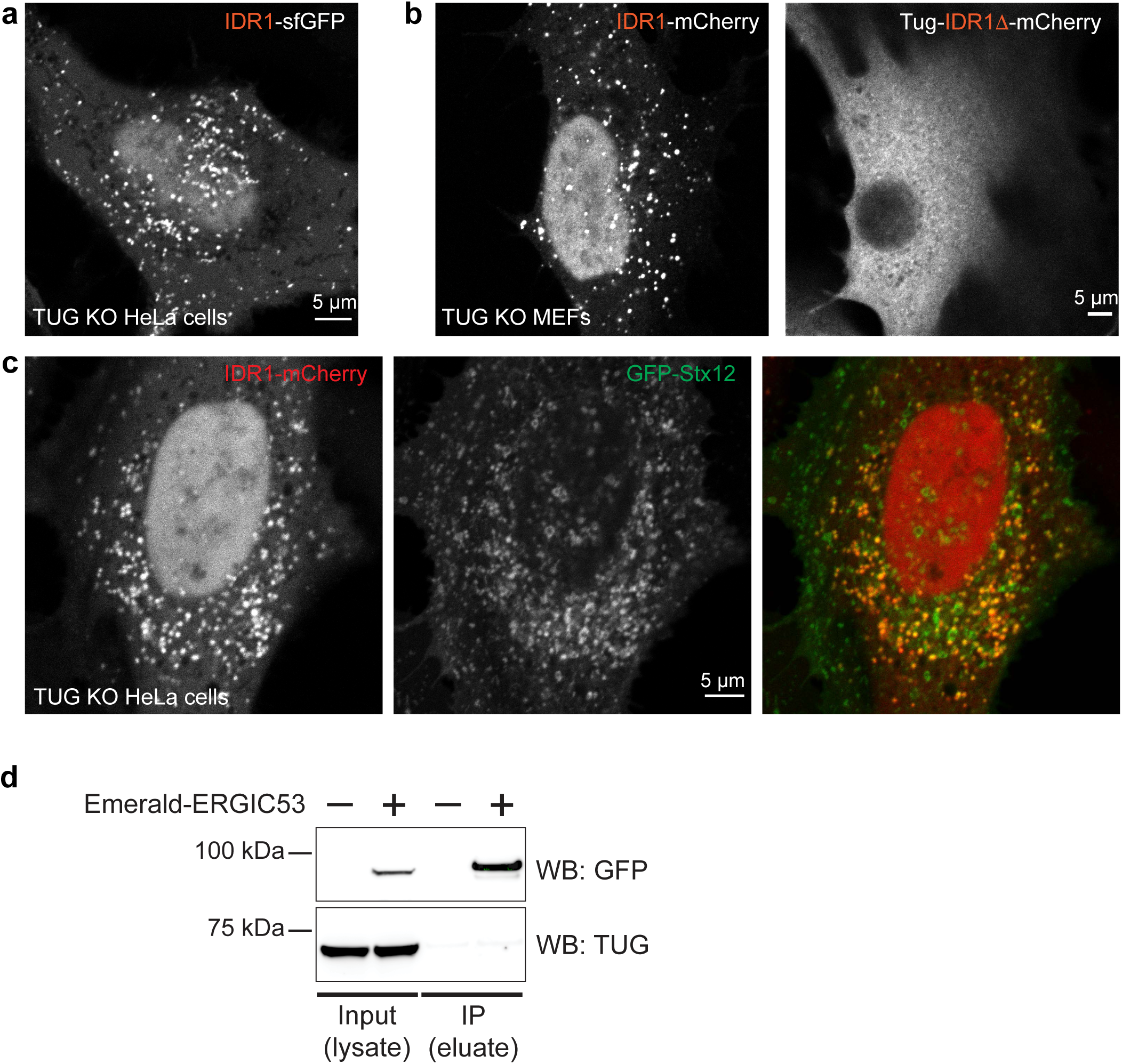
a) Images from TUG KO HeLa cells transfected with IDR1 tagged with monomeric superfolder GFP (IDR1-sfGFP) show that IDR1 distribution is similar to that observed upon expression of the IDR1-mCherry construct (see Fig. 3b). This indicates that the distribution of the IDR1 in cells is independent of the fluorescent tag or the linker sequence, and that it is inherent in the protein sequence. b) Images of TUG KO MEFs infected with retroviruses to express IDR1-mCherry (left) or Tug-IDR1Δ-mCherry (right). Cells were images 48 hours after infection. Similar to the distribution of these proteins in TUG KO HeLa cells (see Fig. 3b, c), expression of IDR1 alone in MEFs results in a punctate distribution, while the deletion of the IDR1 results in a diffuse distribution and exclusion of the fluorescent signal from the nucleus. c) Images from TUG KO HeLa cells co-transfected with IDR1-mCherry and GFP-Stx12. Similar to the full-length protein (see Fig. S1e), IDR1 puncta also show colocalization with a subset of GFP-Stx12 structures. d) HEK293T cells were transfected with Em-ERGIC53, and the protein was immunoprecipitated using GFP trap agarose beads. In the western blots, cell lysates (input) and eluates from the immunoprecipitated fractions were probed using the GFP antibody (top) and TUG antibody (bottom). Bands corresponding to Em-ERGIC53 are seen in cell lysates and in IP eluates obtained from cells transfected with this plasmid, and this protein is absent from the controls, indicating specificity in the immunoprecipitation. The TUG protein band is only seen in the input fractions, and is not co-immunoprecipitated with ERGIC53 using these experimental conditions.

